# Wing shape evolution is not constrained by ancestral genetic covariances in the invasive *Drosophila suzukii*

**DOI:** 10.1101/2024.01.02.573869

**Authors:** Antoine Fraimout, Stéphane Chantepie, Nicolas Navarro, Céline Teplitsky, Vincent Debat

**Affiliations:** Institut de Systématique, Evolution, Biodiversité, ISYEB – CNRS, MNHN, UPMC, EPHE, Université des Antilles, Muséum National d’Histoire Naturelle, Sorbonne Universités, 57 rue Cuvier, 75005 Paris, France; UMR CNRS 6282 Biogéosciences, Université de Bourgogne-Franche-Comté, 21000 Dijon, France; Centre d’Ecologie Fonctionnelle et Evolutive, CEFE – UMR 5175 – CNRS, IRD, EPHE, Université de Montpellier, Université Paul Valéry, 34293 Montpellier, France

**Keywords:** *Drosophila suzukii*, **G**-matrix, geometric morphometrics, biological invasion, quantitative genetics, Q_ST_-F_ST_

## Abstract

The extent to which phenotypic evolution can be constrained by genetic correlations is an important question in evolutionary biology. To address this question, biological invasions are opportune models where derived, invasive populations can be compared to their extant ancestors, allowing to track the evolution of genetic correlations from the ancestor, throughout the invasion process. In this paper, we focused on the worldwide invasion of *Drosophila suzukii* (Matsumara, 1931), and investigated the evolution of the genetic covariance matrix **G** of wing shape between ancestral native, and derived invasive populations. Leveraging demographic history resolved by population genetics approaches, we tested whether **G** remained stable during the invasion. Using a multivariate Q_ST_-F_ST_ approach, we further tested whether or not the observed phenotypic divergence in wing shape aligned with a neutral scenario of evolution. Our results show moderate yet significant quantitative genetic differentiation of wing shape among *D. suzukii* populations and a relative stability in the structure of **G**, presenting a roughly spherical shape but slightly different volumes. These characteristics likely reflect the demographic history of populations and suggest a low level of genetic constraint on wing shape evolution. The divergence between populations was greater than expected under a purely neutral model of evolution, compatible with an effect of divergent selection among them. Overall, our study suggests that selection and drift, but not ancestral genetic constraints, affected the early stages of wing shape evolution during *D. suzukii* invasion.

## Introduction

Biological invasions often involve stochastic demographic changes through founder effects and genetic bottlenecks (Golani *et al*., 2007) or the shuffling of different alleles by genetic admixture between populations (Rius & Darling, 2014). Invasive populations can also be subjected to multiple and often novel selective pressures in the new environment and throughout the invasion process (Kilkenny & Galloway, 2013). Consequently, they tend to show marked genetic and phenotypic differences compared to their source populations, and distinguishing whether this differentiation results from adaptive or neutral evolution is challenging (Lee, 2002; Wares, 2005; Facon *et al*., 2006; Hayes & Barry, 2008; Keller & Taylor, 2008; Prentis *et al*., 2008; Bock *et al*., 2015; Dlugosch *et al*., 2015; Estoup *et al*., 2016). Furthermore, these evolutionary changes can happen over relatively short time scales (Colautti & Lau, 2015). Hence, a topical question in invasion biology is how fast can invasive populations evolve in their new environment? Can we detect adaptive evolution occurring at the onset of the invasion (Szűcs *et al*., 2017)?

In a new environment, the evolution of a population’s mean phenotype towards a new fitness optimum (*i.e.* adaptation) is conditioned by the genetic variances and covariances of the traits under selection, summarized in **G**, the genetic variance-covariance matrix (Lande, 1976). For a set of traits measured in a given population, the **G** matrix informs about genetic variation for each individual trait (diagonal elements) and for the genetic covariation between traits (off-diagonal elements). Statistical analysis of **G** through an eigen decomposition can then identify the major axis of variation in the population, corresponding to the combination of traits with the most genetic variation, also known as **g**max (Schluter 1996). Identifying the axes of genetic covariance between traits is particularly important as theory predicts that they will influence the evolutionary response: adaptation will be facilitated if selection is aligned with **g**max, or constrained if it is in a direction where there is little, or no genetic variation for the combination of traits under selection (Arnold *et al*., 2001, Agrawal & Stinchcombe, 2009; Fischer *et al*., 2016). This is formalized in the multivariate breeder’s equation (Lande, 1979):

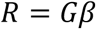

where the response to selection *R* - the change in average phenotype of the population in one generation - equals the product of **G** and the directional selection gradient *β*.

It has been proposed in the context of evolutionary diversification, that the **G** matrix of an ancestral population can constrain derived populations to diverge along certain directions of the phenotypic space (*i.e.* the lines of least evolutionary resistance; Schluter, 1996). For instance, early stages of adaptive phenotypic divergence could be constrained by the direction of the major axis of the ancestral **G** matrix (*i.e.* ancestral **g**max; Schluter, 1996; McGuigan, 2006; Walter *et al*., 2018) and result in an alignment between **g**max and the leading vector of divergence of the among-populations divergence matrix (**D**). On the other hand, under a neutral model of phenotypic evolution, divergence among populations is also expected to take place along that same axis of greatest genetic variation (Lande 1979) and result in proportional **D** and **G** matrices (Chenoweth *et al*. 2010). In other words, a concordance between the major axes of **D** and **G** could be both suggestive of neutral evolution (Arnold *et al*. 2001; Bégin & Roff, 2003) or indicative of selection.

Furthermore, valid predictions of evolutionary trajectories depend on whether or not **G** remains stable across space and time (Jones *et al*., 2004; Arnold *et al*., 2008). The stability of **G** has received much attention (*e.g.* Roff, 2000; Steppan *et al*., 2002; Bégin & Roff, 2003; Mezey & Houle, 2003; Björklund, 2004; Blows & Hoffmann, 2005; Arnold *et al*., 2008; Walsh & Blows, 2009) and its sensitivity to selection, migration and drift has been investigated both theoretically (*e.g.* Jones *et al*., 2004; Guillaume & Whitlock, 2007, Chantepie & Chevin, 2020), and empirically under natural and laboratory conditions (Merilä *et al*., 1994; Phillips *et al*., 2001; Conner *et al*., 2003; Cano *et al*., 2004; Eroukhmanoff. & Svensson, 2011; Björklund *et al*., 2013; McGlothlin *et al*., 2018; Walter 2023; Henry & Stinchcombe 2023; Chantepie *et al*. 2024) but no clear consensus has emerged. For instance, the recent work of Henry & Stinchcombe (2023) has shown that the **G** matrix of floral traits in the ivy (*Ipomoea hederacea*) remained stable across a large latitudinal gradient in continental USA, despite evidence of phenotypic divergence among populations. Conversely, Mallard *et al*. (2023) have shown using a large experimental evolution study that the orientation of the **G** matrix of locomotor traits in *C. elegans* can be altered by genetic drift, despite phenotypic stasis. Multiple additional lines of evidence suggest that **G** can be reshaped by selection (Jones *et al*., 2003; Jones *et al*., 2004; Dugand *et al*. 2021), drift (Phillips *et al*., 2001) and environmental conditions (*e.g.* Sikkink *et al*., 2015; Wood & Brodie, 2015) even over the span of a single generation (Siren *et al*., 2017), but, there is also recent evidence for stable **G** matrices across populations and continents (*e.g*., Puentes *et al*., 2016; Delahaie *et al*., 2017, Henry & Stinchcombe 2023). Overall clear-cut predictions on the interplay between **G** and phenotypic divergence in response to selection or drift are difficult to formulate, particularly at a contemporary timescale. Therefore, the stability of **G** needs to be further explored empirically along with clear testing of the neutral expectation for among-population divergence against observed data (Merilä & Björklund 2004).

Here, we focus on the worldwide invasion of the spotted wing drosophila, *Drosophila suzukii* (Matsumura, 1931) to assess the stability of **G** between native and invasive populations across continents. *Drosophila suzukii* is a highly invasive pest of agricultural crops in Europe and North America where it was first detected almost simultaneously in 2008 (see Asplen *et al*., 2015 for review). We reconstructed the detailed history of this worldwide invasion and showed the presence of multiple independent introductions from Asia involving population bottlenecks and genetic admixture events within invaded regions (Fraimout *et al*., 2017). A consequence of this demographic history is that native and invasive populations show significant levels of neutral genetic differentiation as measured by F_ST_ estimated using microsatellite loci (Fraimout *et al*., 2015; Fraimout *et al*., 2017). At the phenotypic level, we showed using laboratory lines derived from samples of natural populations that native (Japanese) and invasive (French and North-American) *D. suzukii* display genetically based differences in wing shape and size (Fraimout *et al*., 2018). However, whether this phenotypic divergence is pervasive across replicated populations, and whether it is associated with changes in the structure of the underlying **G** matrix is unknown. This invasion therefore provides an opportune context to address long-standing questions about the evolution of multivariate phenotypes and the structure of genetic variances and covariances in colonizing populations. Particularly the presence of multiple invasive populations within Europe and USA allows assessing the magnitude and direction of phenotypic evolution from the native ancestral populations. In other words, it is possible to estimate how much the mean phenotype of each invasive population has diverged from the ancestral (*i.e.* native) population and whether phenotypic evolution repeatedly occurred in the same direction or at random, in all directions of the phenotypic space. Given the availability of extant ancestral populations, this in turns allows to determine whether phenotypic divergence among invasive populations has been constrained by ancestral genetic covariances. Furthermore, comparing populations with known history could help identify the effects of demographic processes on the structure of the **G** matrix. For instance, we expect that periods of bottlenecks or population admixture – all experienced by invasive *D. suzukii* populations to a different extent (Fraimout *et al*., 2017, Olazcuaga *et al*. 2020) – may have altered the structure of **G** in invasive populations. Specifically, depletion of genetic variance due to bottlenecks could result in a **G** matrix of smaller volume. Alternatively, new combinations of alleles produced by genetic admixture, could increase the volume or **G**, or alter its shape, by changing the distribution of genetic variance over the first axes of variation. Our main objective was thus to investigate whether or not **G** remained stable throughout the invasion history of *D. suzukii*. Additionally, access to both molecular and quantitative measures of genetic variation allows testing for a signal of selection in the observed phenotypic divergence. By comparing F_ST_ calculated from molecular markers to its phenotypic equivalent Q_ST_, it is possible to determine whether phenotypic divergence exceeds or is equal to the divergence expected under genetic drift alone (Spitze 1993, Whitlock 2008). While the former (Q_ST_ > F_ST_) would be indicative of divergent selection acting on the study traits, the latter (Q_ST_ = F_ST_) would suggest neutral phenotypic evolution.

We used an isofemale line breeding design (Hoffmann & Parsons, 1988; David *et al*., 2005; Berger *et al*., 2013) and a geometric morphometric approach (*e.g.* Klingenberg 2003) to estimate the quantitative genetic divergence in wing shape between six natural populations of *D. suzukii* sampled in the native area (Japan) and the two main invaded areas in Europe and the USA. We specifically addressed the following questions: i) how did wing shape evolve across multiple invasive populations? ii) did the **G**-matrix remain stable during the invasion**?** iii) has phenotypic divergence among invasive populations been constrained by ancestral genetic covariances and iv) can it be attributed to selection acting on wing shape?

## Material and methods

### Samples, lines and data acquisition

*Drosophila suzukii* adults were sampled in 2014 using banana bait traps and net swiping at six localities representative of their global distribution range in Japan (native range), USA and France (invasive range) (Fig. 1). Within each country, we sampled two localities representative of a northern and a southern latitude. Repeated clinal evolution of wing morphology has been described in the related invasive species *Drosophila subobscura* and proposed as an adaptation to different temperature optima at northern and southern latitudes (Huey *et al*., 2000; Gilchrist *et al*., 2001; Gilchrist *et al*., 2004). Thus, we hypothesized that sampling populations close to the extremes of their latitudinal distribution would maximize our chance to capture possibly adaptive phenotypic divergence in the invaded range. In USA and France, these localities were also representative of the invasion history, with the southern localities corresponding to the first reported occurrences of *D. suzukii* (Watsonville, USA and Montpellier, France; Hauser, 2011; Calabria *et al*., 2012) and the northern localities representing the expansion front of the species in the area at the time of sampling (Dayton, Oregon, USA and Paris, France). It is worth stressing that although the most likely genetic source of Europe and USA invasions was identified as continental Asia (*i.e.* China, Fraimout *et al*., 2017), *F* statistics and Bayesian clustering methods indicate weak genetic differentiation between these sources and our Japanese population samples, thereby suggesting that the use of Japanese populations as ancestral representatives is an acceptable approximation. For the Japanese flies, a northern population was sampled in Sapporo (Hokkaido Island) and a southern population was sampled in Tokyo (Honshu Island). In both Japanese localities, traps were placed in gardens inside the university campuses. In the invasive North American range, sampling habitats corresponded to vineyards in Dayton, and strawberry fields in Watsonville. French samples were collected in the botanical garden of the Natural History Museum in Paris and close to vineyards in Montpellier. 30 isofemale lines on average (see Table S1 for details and population codes) were established for each locality by placing randomly chosen wild-caught gravid females in separate rearing vials, and splitting F_1_ offsprings in five replicate vials. Following the isofemale approach, we further expanded each line by letting individuals mate freely within tubes (*i.e*., full-sib mating) and transferring their progeny to new rearing vials each generation. We repeated this for five generations and in the fifth generation, a single female per line was isolated in a new rearing tube to produce the focal offspring used in the study. At the end of the experiment, 5 to 10 full-sib males were randomly chosen from each line from each population for phenotyping. Rearing medium consisted of cornstarch and yeast with added anti-biotic (hydroxy-4 benzoate). Lines were randomly positioned in incubators and kept for the five generations at 22°C. A total of 1323 individuals distributed across 177 isofemale lines were phenotyped. Wings were carefully clipped from over-anesthetized flies using tweezers and mounted on glass slides in a mixture of ethanol and glycerin. Coverslips were sealed using nail polish and maintained with small weights to flatten the wing as much as possible. Images of the wings were acquired with a Leica DFC 420 digital camera mounted on a Leica Z6 APO microscope. Fifteen landmarks were defined on the dorsal side of the right wing and digitized using a custom *ImageJ* plugin written by C. P. Klingenberg (pers. comm.).

**Figure 1.**
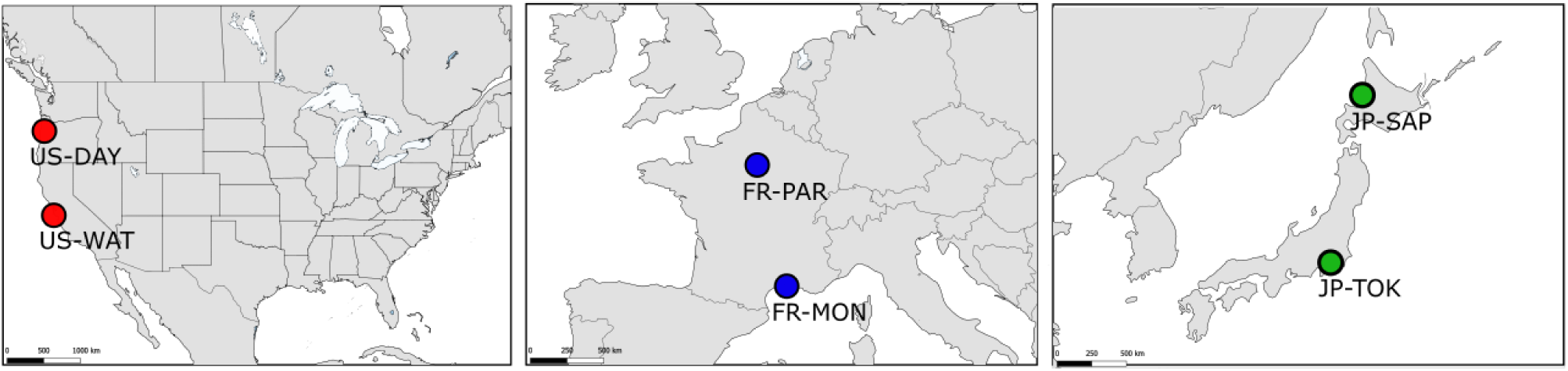
Sampling locations of the six *D. suzukii* populations. Adult *D. suzukii* were sampled from the wild in 3 countries (dark grey shading) from the native range in Japan (green circles) and the invasive range in France (blue circles) and the USA (red circles).

### Wing shape differentiation – Geometric morphometric analyses

We focused our study on the analysis of wing shape variation among *D. suzukii* populations. Shape data have been used in multiple quantitative genetics studies (Klingenberg & Leamy, 2001; Klingenberg, 2003; Monteiro *et al*., 2003; Hansen & Houle 2004; Mezey & Houle, 2005; Myers *et al*., 2006; Polly, 2008; Klingenberg, 2010; Adams, 2011; Leinonen *et al*., 2011; Martínez-Abadías *et al*., 2012) and are suitable for the estimation of **G**, and relevant for the study of multivariate phenotypic evolution. We followed a standard geometric morphometrics protocol (Bookstein 1996) to characterize wing shape: first, Procrustes generalized least squares superimposition was used to extract shape information from the landmarks coordinates (GLS, *e.g.* Dryden & Mardia, 1993). Briefly, this method allows to scale, rotate and translate the configuration of landmarks to obtain shape information that is free of orientation or size differences. Second, a principal component analysis (PCA) was applied to the set of 30 coordinates (15 landmarks in 2 dimensions x and y) obtained after superimposition. From the PCA, 26 non-null components were conserved as input data (corresponding to the 30 coordinates minus 4 degrees of freedom lost through the scaling, rotation and translations (x and y)). We estimated the centroid size of the wing as the square root of the sum of the squared distances of all landmarks from their centroid, and used this centroid size as a size variable in the subsequent analyses. As divergence in wing shape first appeared to be weak on the PCA plot, we further performed a between-group principal component analysis (*bgPCA*) to visualize the overall variation in wing shape among populations (*e.g.* Mitteroecker & Bookstein, 2011). We then estimated the statistical significance of the shape divergence among populations by applying a multivariate analysis of variance (MANOVA) using the 26 PCs as response variables, and population of origin as a fixed effect. We further estimated the pairwise differences among populations with permutational pairwise MANOVAs, using the *RVAideMemoire* R package (v.0.9-55, Hervé, 2016). We used the Wilk’s lambda statistics and set the number of permutations to 10000 while correcting for multiple testing using the *fdr* method (Benjamini & Hochberg 1995). Finally, to account for any allometric effect (i.e. a change in shape associated with a change in size,e.g. Klingenberg 2010), we also fitted the MANOVAs using the residuals of a size/shape multivariate regression, corresponding to the allometry-free components of wing shape variation.

### Wing shape differentiation – phenotypic divergence vector analysis

To estimate the divergence in wing shape among populations of *D. suzukii*, particularly the magnitude and direction of phenotypic evolution in the derived invasive populations from their Asian ancestors, we applied a geometric approach to phenotypic evolution (*e.g.,* Stuart *et al*. 2017, Fraimout *et al*. 2022), by calculating vectors of phenotypic divergence among populations. The phenotypic mean of each population was first estimated from the MANOVA (see section above) using the 26 non-null PC representative of wing shape, and accounting for differences in wing size by adding the wing centroid size as covariate. Then, we estimated the differences in mean wing shape between each invasive population and the ancestral wing shape, assessed as the average of the two Japanese populations (Fig. 2). We also estimated divergence at the country level, using the three countries average wing shapes. In total, we obtained six phenotypic divergence vectors describing the difference between the ancestral Japanese and each invasive population (4 vectors) and between the ancestor and each country’s mean wing shape (2 vectors). To test whether phenotypic divergence occurred along similar or different directions, we assessed the similarity between these vectors of divergence by comparing their angle to the distribution of angles between 10,000 pairs of random vectors of the same dimensionality (Klingenberg and McIntyre, 1998). Angles between vectors were calculated as:

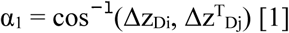

**Figure 2.**
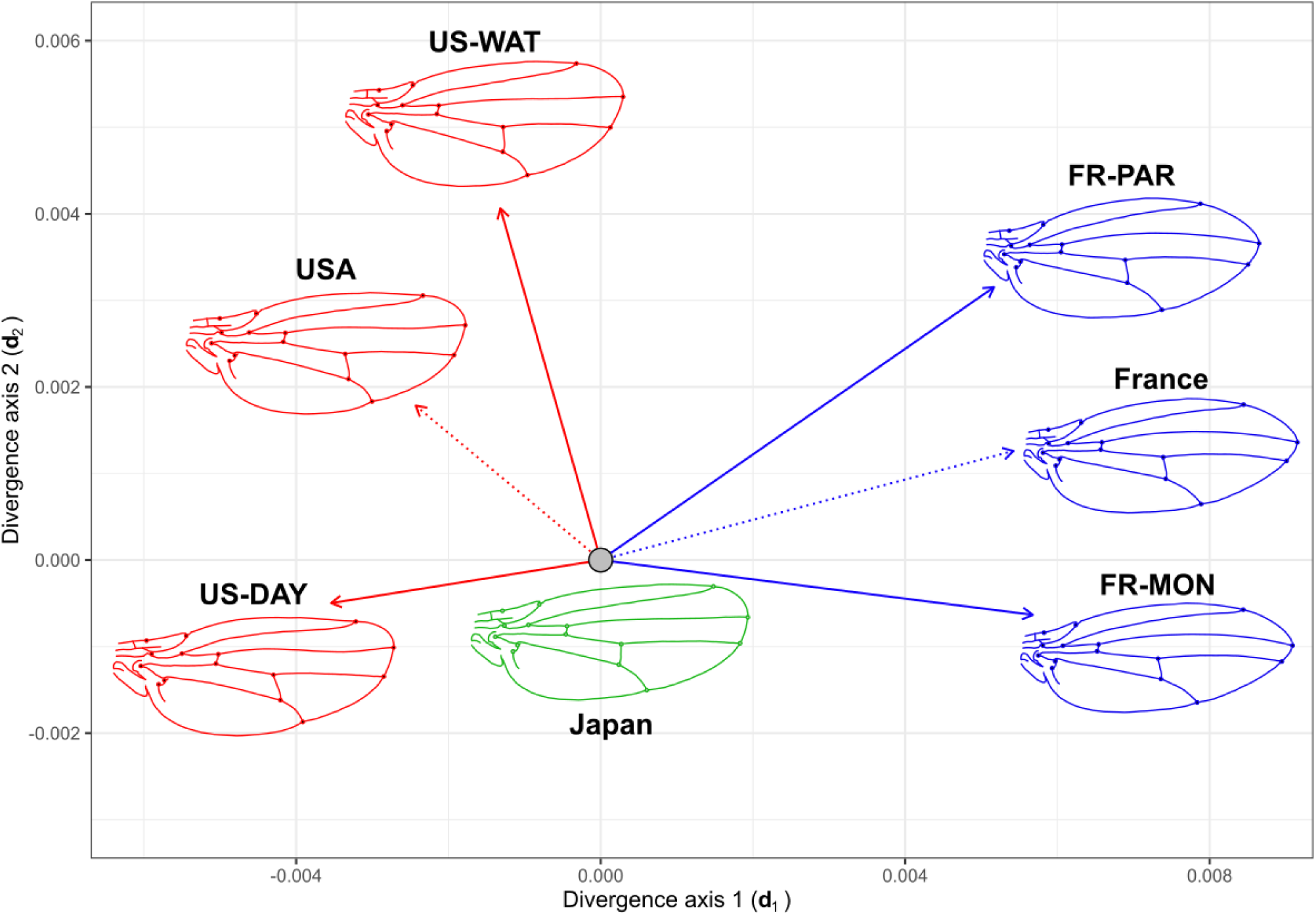
Phenotypic divergence of the invasive populations from their common ancestor. Graphical representation of the vectors of phenotypic divergence of each invasive population (solid line) from their hypothetical common ancestor (Japan, grey filled circle). The vectors are projected on the multivariate divergence space where d1 and d2 represent the first and second main axes of the multivariate divergent covariance matrix. The magnitude (length of the arrow) and direction (angles between arrows) of the vectors reflect the difference in mean wing shape phenotype among groups. Dashed lines represent the vector of phenotypic divergence of the average wing shape by country (USA or France) from the Japanese hypothetical ancestor. Colors correspond to Japanese (green), French (blue) and American (red) wing shape. Wing shape outlines represent the mean shape of each group and show the wing shape evolution from the Japanese ancestor. Outlines were obtained via a thin plate spine warping of a reference outline using MorphoJ (Klingenberg, 2011), based on the position of landmarks and are used for visualization purposes only.

where z_D_ corresponds to a divergence vector of population *i* or *j* from the ancestor. An eigenanalysis of the matrix containing the four population vectors was applied to synthesize the main axes of divergences.

### Matrices estimations – population-specific **G** matrices

We estimated the six **G** matrices corresponding to the six sampled populations under a Bayesian framework using the R-package *MCMCglmm* (Hadfield, 2010; Fig. 3). To this end, we partitioned the phenotypic variance in wing shape for each population using a multivariate animal model (Kruuk, 2004) of the form:

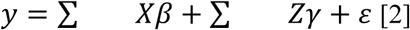

**Figure 3.**
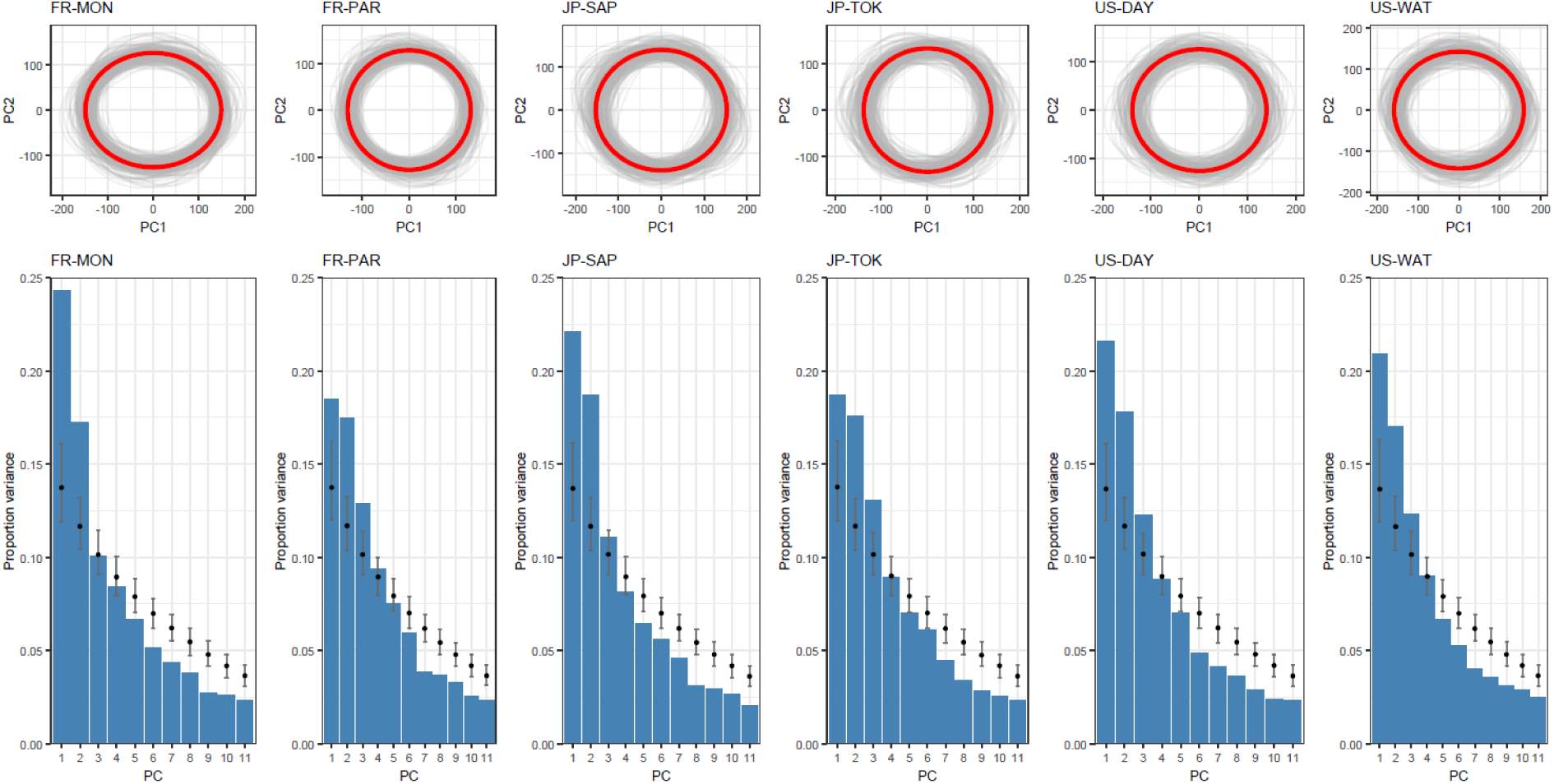
Graphical representation of the population-specific G-matrices. For each population-specific **G** matrix, the projection of the mean **G** matrix (red circles) onto the first two axes of the eigen decomposition (PC1 & PC2) are shown (top 6 panels). For each population, a sample of 200 ellipses corresponding to 200 random posteriors drawn from the MCMC outputs (grey ellipses) illustrate the uncertainty around our estimated **G** matrices. Results from the eigen decomposition of each matrix is shown by plotting the percentage of variance explained by the first 11 PCs (blue histograms; mean total variance explained across PC and populations = 91.22%) for both observed matrices, and null matrices (black filled points). This shows that the main axes of variation (PC1 and PC2) explain more variance than expected at random. It is important to note that no conclusions should be drawn beyond PC3, as differences in eigenvectors between null and observed **G** matrix prevent interpretation.

where *y* is the multivariate vector of phenotypic values for wing shape corresponding to the 26 non-null principal components (PCs) accounting for the totality of shape variance; *β* is the vector of fixed effect; γ is the vector of random effects; ε is the vector of residual errors and *X* and *Z* the design matrices relating to the fixed and random effects, respectively. We estimated **G** matrices separately for each population using model [2] and appended the matrix of relatedness between individuals to the vector of random effects with the *pedigree* option implemented in the *MCMCglmm* function. The pedigree corresponding to our crossing design was coded in R using custom scripts (see *Data availability* statement). In all models, wing centroid size was fitted as a fixed effect. We used a prior set with variance estimates following a ‘broken-stick’ model of decrease of variance, appropriate for the use of PC scores representing decreasing proportion of explained variance (Jolliffe, 2002), with null covariances and a degree of belief *nu* = n + 0.002 (where n is the number of traits, Houle & Meyer, 2015). We ran each model for 1 200 000 iterations with a burnin period of 200 000 and a thinning rate of 1000, resulting in 1000 Monte Carlo Markov Chain (MCMC) posterior samples. The convergence of models was checked visually from plots of MCMC chains and posterior distributions as well as autocorrelation level (<0.05 in all models). All analyses were performed in R (v.4.2.2; R Core Team, 2022).

### Test of **G**-matrix stability – Differences in the size and shape of **G**

One of our main objectives was to assess the stability of the **G** matrix in the face of the demographic changes linked to the invasion. According to previous results (Fraimout *et al*. 2017) all invasive *D. suzukii* populations have experienced genetic bottlenecks, and some populations show evidence of genetic admixture from differentiated sources. Our hypothesis was that the variation in total genetic variance due to these demographic events - namely, reduction of genetic variance by bottlenecks or increased genetic variance from admixture - should be reflected in the structure of **G**. To test this hypothesis, we estimated three parameters pertaining to the structure of **G** and performed pairwise comparisons among all six studied populations. Matrix size (*i.e*., volume), estimating the overall amount of genetic variance for wing shape in the population, was calculated as the sum of the matrix eigenvalues. Matrix shape, estimating whether some directions of wing shape variation present more genetic variance than others, was assessed by the matrix eccentricity (*i.e.* the ratio of the first to the second eigenvalue; Jones *et al*., 2003). Matrix shape was also estimated by the proportion of variance along **g**_max_. Significance of metric differences between populations was assessed by estimating the ratio between each population metrics and examining the overlap between the 95% highest posterior density (HPD) of these ratios calculated from the MCMC posterior samples, with zero.

### Test of **G**-matrix stability – Genetic covariance tensor

We further tested for differences among **G** matrices by following the approach of Hine *et al*. (2009) and Aguirre *et al*. (2014) using a fourth-order genetic covariance tensor analysis (hereafter ‘covariance tensors’). Briefly (see details in Hine *et al*. 2009; Aguirre *et al*. 2014), variances and covariances among all elements of the original set of **G** matrices are calculated to construct the fourth-order covariance tensor **Σ** and the second-order (*i.e.* matrix) tensor **S**. Eigenanalysis of the **S**-matrix allows for the construction of eigentensors which represent independent vectors of differences in variance and covariance among **G** matrices (Fig. 4). Covariance tensors provide a way to compare more than two genetic covariance matrices and describe independent axes of divergence among them. In other words, covariance tensors allow identifying trait combinations explaining most genetic variation among **G** matrices. We applied the covariance tensor analysis to our set of six **G** matrices using the full 1000 MCMC samples for each matrix and compared the observed divergence to the null expectation obtained from the analyses of random **G** matrices. We used an approach based on Morrissey *et al*. (2019) to simulate the null distribution of **G** matrices. For each population, the breeding values of founders were drawn from a multivariate distribution with zero mean and a variance-covariance matrix equal to the **G**-matrix estimated from this population with the animal model [2]. Breeding values were then randomly shuffled among the founders and the breeding values of all remaining individuals were estimated based on the pedigree of each population. As our breeding design included five generations of inbreeding, we modeled a pedigree to better approximate the actual relatedness of our measured individuals. The breeding values of an individual were drawn from a multivariate normal distribution with a vector of means equal to the mean breeding values of its parents and a deviation equal to the segregation variance (Walsh & Lynch, 2018). For an individual, the segregation variance was estimated as half the product of the randomized **G** matrix of their population (estimated on the new population’s founders) and one minus the average inbreeding coefficient of its parents (Walsh & Lynch, 2018, pp.557). The phenotypes were then recomposed by adding the random and fixed effects estimated by the animal model following [2].

**Figure 4.**
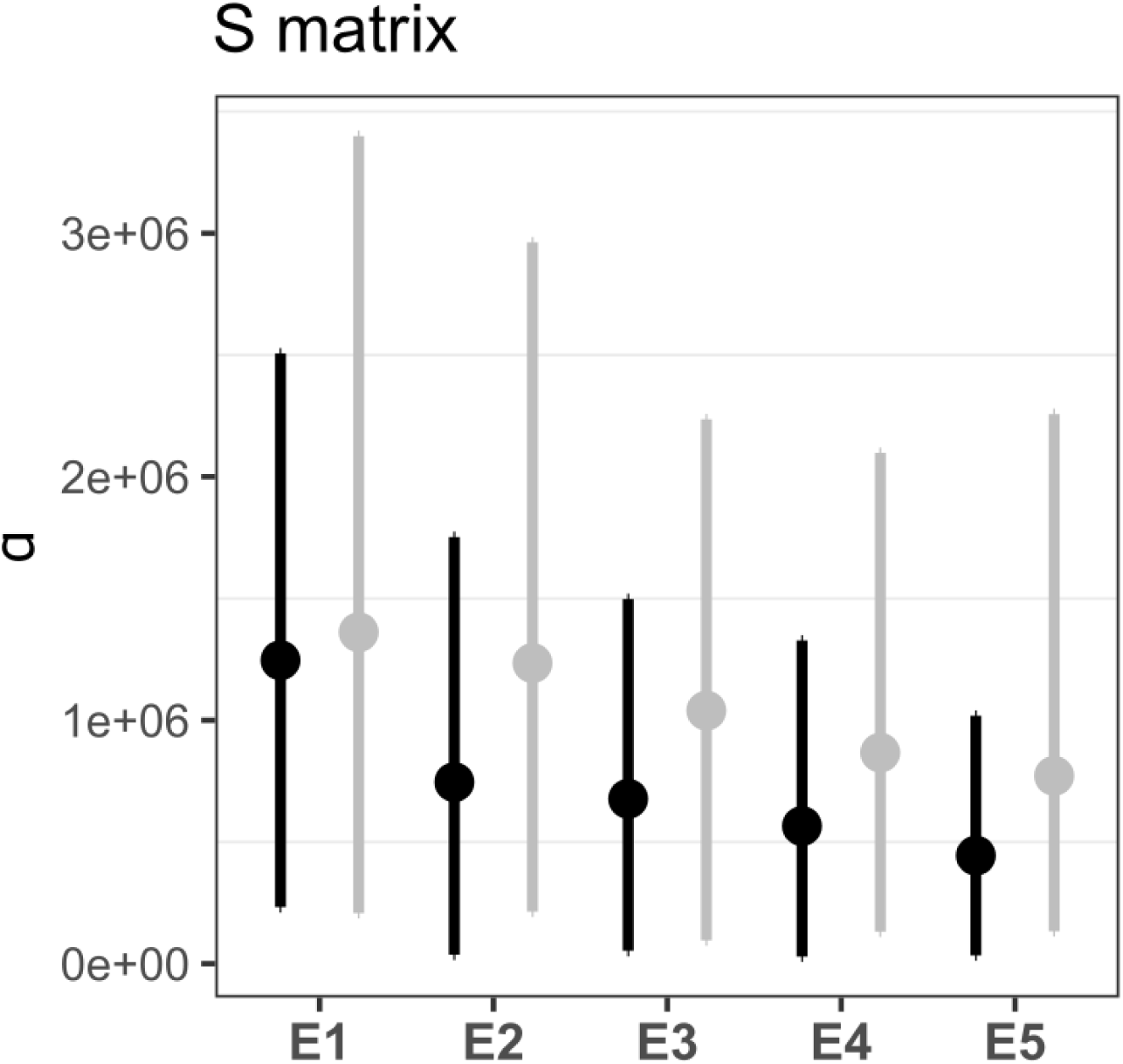
Results of the genetic covariance tensor analysis. Eigenvalues of the 5 observed eigentensors of **S** (black) are shown against the random expectation from the estimated null **G** matrices(grey). Overlapping of the 95% confidence intervals indicates that the six observed **G** matrices are not more different than a random set of **G** matrices.

We repeated this sequence for each posterior sample of animal models and resulted in 1000 null datasets for each population. The **G** matrix of null datasets was estimated following eq. [2]. To reduce computational burden, we sampled 200 posterior samples for each null animal model (Morrissey *et al*., 2019) to compute a mean null distribution **G** matrix for each population that accounts for both the sampling effect and the uncertainty in the **G** matrix estimates. Each final null **G** matrix consisted of the average of the 1000 MCMC posterior for each of the 100 replicates and were used to estimate random genetic covariance tensor as described in Aguirre *et al*. (2014) using custom scripts in R (see *Data availability* section for source code),

### Test of genetic constraints: proportionality between **G** and **D**

In the case of populations deriving from a common ancestor, quantitative genetics theory predicts that genetic constraints imposed by the ancestral **G**-matrix can influence among-population divergence in a manner, leading to the proportionality of the divergence matrix **D** and the ancestral **G** under neutral or adaptive scenarios (Lande, 1979; Merilä & Björklund 1994, 2004). This proportionality induced by genetic constraints is expected under selection, where the evolutionary response is biased along the ancestral main directions of genetic variation (e.g. Schluter 1996, McGlothlin et al 2018), but also under neutral evolution, as genetic drift is expected to induce a global reduction in genetic variances, but no predictable change in the orientation of the divergence, leaving on average **G** and **D** proportional (Lande 1979; Phillips *et al* 2001).

The similarity between **D** and **G** was first investigated by computing the angle between a divergence vector ***d*** and the ancestral **G** matrix following Krzanowski (1979):

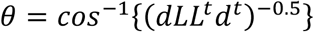

where **L** is the set of first *q* eigenvectors of **G**, where *q* was set equal to half the shape space dimensionality (*q* = (2*k* – 4)/2 = 13; and *k* is the number of digitized landmarks) and ***d*** is a divergence vector..

We compared the observed angles to their random expectations by simulating 1000 random vectors *v*_2*k*_ projected in the tangent space according to

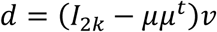

where ***µ*** is the vectorised mean shape (Dryden and Mardia, 1998).

### Test of selection: multivariate Q_ST_-F_ST_ comparison

A proportionality between **G** and **D** is indicative of an effect of genetic constraints on phenotypic evolution, but is insufficient to identify the evolutionary process at play, pointing at the need for a more stringent test to decipher the adaptive and neutral scenarios (Martin *et al*. 2008). To test the hypothesis that phenotypic evolution among invasive *D. suzukii* populations resulted from divergent selection on wing shape, we used the two steps method proposed by Martin *et al*. (2008). Briefly, this method is a multivariate extension of the classical Q_ST_-F_ST_, testing whether the observed phenotypic divergence is greater (Q_ST_ > F_ST_), equal to (Q_ST_ = F_ST_), or smaller (Q_ST_ < F_ST_) than expected under drift alone (see Martin *et al*. 2008 and Chapuis *et al*. 2008 for more details). First, proportionality between **D** and **G** was tested using the MANOVA estimates of the covariance matrices. Second, the coefficient of proportionality *ρ_ST_* between the ancestral **G** matrix and the divergence matrix **D** was estimated and compared to the predicted neutral divergence 2F_ST_/(1-F_ST_) obtained from neutral molecular markers (Rogers & Harpending 1983; Martin *et al*. 2008). While *ρ_ST_* is not strictly equivalent to Q_ST_, the rationale behind its comparison to a measure of neutral differentiation (*i.e*. F_ST_) is very similar. ρ_ST_ is expressed as 1/n_f_(ρ_MS_ - 1) (equation 11 in Martin *et al*. 2018; pp. 2139), where n_f_ is the number of families sampled, and ρ_MS_ is the coefficient of proportionality between the among- and within-population mean square matrices, estimated from phenotypic data. Comparing the confidence intervals of ρ_ST_ to that of 2F_ST_/(1-F_ST_) estimated from molecular markers provides a test of neutrality similar in rationale to the Q_ST_-F_ST_ comparison: in case of an overlap between the CIs, the neutrality hypothesis is not rejected, this is similar to observing Q_ST_ ≈ F_ST_. Conversely, deviation from this overlap would be indicative of selection. To test for proportionality between **D** and **G**, we used the Bartlett-adjusted test of proportionality between matrices as recommended in Martin *et al*. (2008). We used the *k.prop* R function of the neutrality package from Martin *et al*. (2008) to calculate *ρ_ST_* and the *hierfstat* R package (v.0.5.11; Goudet & Jombart 2015) to get bootstrapped estimates of F_ST_ from 25 neutral microsatellite loci (Fraimout *et al*. 2015, 2017). Inputs for both tests were obtained by performing a MANOVA on the wing shape data and using the log-centroid size as covariate and population of origin as fixed effect. Also following recommendation from Martin *et al*. (2008), we used the first 11 PCs describing wing shape variation to maximize accuracy of the statistical tests so that *p <* 2*nb* (where *p* is the number of traits and *nb* is the number of populations sampled) while retaining PCs accounting for > 90% of variance.

### Test of selection: solving the multivariate breeder’s equation

It has been suggested that in Drosophila, flight performance in different thermal conditions might be under selection, favouring the evolution of latitudinal gradients in wing size and shape (e.g. Gilchrist *et al*. 2001). This hypothesis is indirectly supported by phenotypic plasticity analyses, showing better flight performance in cold air in flies reared at low temperatures, associated with different wing morphologies (Frazier *et al*. 2008). While we are mostly oblivious to the selective factors affecting *D. suzukii* invasive success (but see Camus *et al*. 2025), and specifically the evolution of its wing morphology, flight performance might play a role in its dispersive ability, or be under selection in the diverse thermal conditions encountered across its latitudinal ranges. We used data obtained in a previous study showing an association between wing shape and flight speed in *D. suzukii* (Fraimout *et al*. 2018), to explore the hypothesis that selection on flight speed might have driven wing shape divergence among invasive populations, using the multivariate breeder’s equation (Lande 1979). First, an experimental selection differential was derived from Fraimout *et al*. (2018) data. In this previous study, we reconstructed 3D flight trajectories of flies reared under different developmental temperatures and measured flight parameters such as speed, acceleration, or sinuosity. Recorded individual wings were digitized and wing landmarks coordinates used to assess the relationship between flight parameters and wing shape (i.e. we identified wing shape variation associated with flight speed variation). Landmark data from this previous study were aligned onto the reference configuration of the present study to allow direct comparability. Solving the breeder’s equation assumes that traits change is correlated with fitness. In the absence of fitness data, we hypothesized that flight performance might significantly contribute to fitness in Drosophila (e.g. Frazier *et al*. 2008) and we therefore used the shape change associated with the strongest flight speed difference as a hypothetical selection differential. Wing shape was regressed onto the log of flight speed using a multivariate regression (*e.g.* Mardia *et al*. 1979; Monteiro 2003) to estimate the vector of shape change associated with highest flight velocity *s*_velocity_. The corresponding response to selection Δ*z* was then predicted using the multivariate breeder’s equation (Lande & Arnold 1983) and the estimated ancestral (*i.e.* Japanese) **G** and **P** matrices:

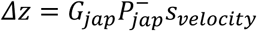

where ^−^ is the Moore-Penrose generalized inverse. This predicted response to velocity selection was compared to the different vectors of population divergences (see above) by computing the angle between them following

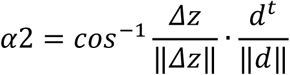

The observed angles were then compared to the expectation of the angle between Δ*z* and 1000 random vectors (see above). Our hypothesis here is that if a high flight speed has been favoured by selection throughout the invasion, we should observe a match between the evolutionary shape changes (the phenotypic divergence among populations) and the shape change resulting from the hypothetical response to selection.

### Test of selection: percentage of genetic variance along divergence vectors

Finally, we reasoned that if selection was involved in the short-term evolution of wing shape, it should manifest in a depletion of genetic variation in the selected directions (*e.g.* Agrawal & Stinchcombe 2009), corresponding to the directions of divergence from the hypothetical common ancestor (Fig. 5). To test this prediction, **G** matrices of each population were projected onto the shape space directions defined by the divergence vectors, *d*^*t*^*Gd*, and expressed in proportion of the total genetic variance. The amount of genetic variance in the direction defined by ***d*** was then compared between the ancestral and derived **G** matrices. We predicted that a reduction in this proportion of variance should be detected if selection was involved in the evolutionary divergence, as depletion of genetic variance due to drift would randomly affect all directions. To test whether genetic variance along the directions of divergence was lower than expected by chance, we compared it to the variance estimated in random directions in the (tangent) shape space.

**Figure 5.**
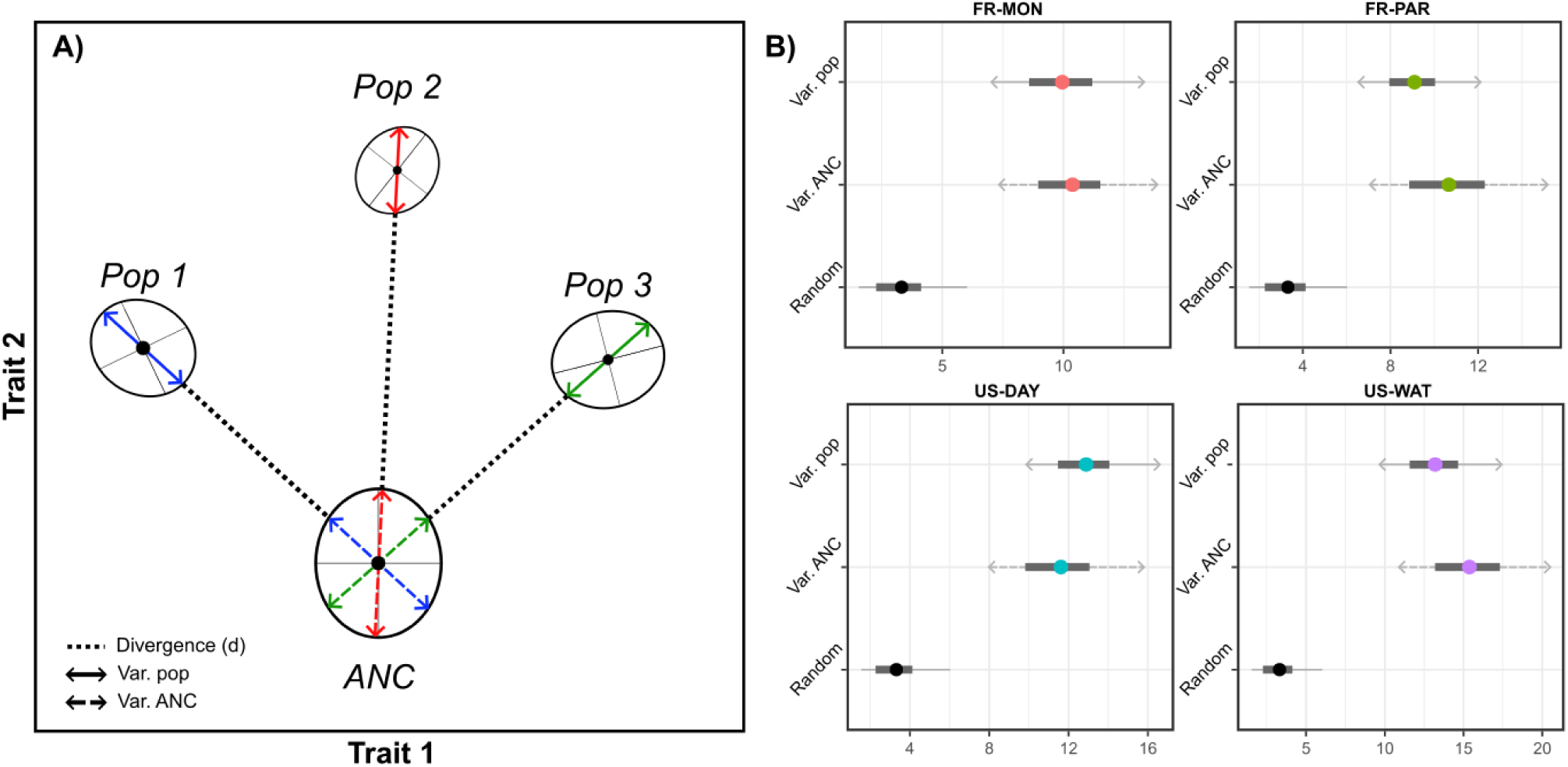
Percentage of genetic variance along divergence vectors. A) Conceptual figure for our test of selection based on the percentage of genetic variance along divergence vectors. In a bivariate trait space, population means (black filled circle) of three populations have diverged from a common ancestor (ANC). The within-population variance (Var.pop) corresponds to the amount of genetic variance in the direction of the phenotypic change between the ancestor (ANC) and the focal population. The ancestral variance (Var.ANC) corresponds to the amount of genetic variance in the ancestral **G** matrix, along that same divergence axis. According to our hypothesis, reduction of the within-population variance (shortening of the Var.pop arrow) would be suggestive of the effect of selection. B) For each population, the observed amount of genetic variation in the direction of divergence was computed (Var. pop.) and compared to the amount of variation in the ancestral **G** matrix in that direction (Var. ANC). The percentage of genetic variance along this axis is plotted (filled grey dot) with the 95% highest posterior density interval (black horizontal lines) next to the population code on the y axis. We hypothesize that selection would result in a reduction in genetic variance in the directions of divergence. This is clearly not observed, as none of the directions shows depleted genetic variance.

## Results

### Significant divergence of wing size and shape across populations

We observed differentiation in wing centroid size between American and all other populations, as well as between the two American populations (Fig. S1): the American populations displayed smaller wing size, with US-WAT displaying the smallest wings. Although the divergence for wing shape among populations was weak, the *bgPCA* showed a signal of geographic differentiation, opposing the pairs of populations according to their geographic origin (Fig. S2). This shape divergence was confirmed by the significant population effect in the MANOVA (Wilk’s lambda = 0.143, df = 5, *p* < 0.001). The MANOVA performed on the residuals of the regression between shape and size also revealed a significant population effect (Wilk’s lambda = 0.199, df = 5, *p* < 0.001), suggesting that the shape difference goes beyond a simple allometric effect. The pairwise permutational MANOVAs revealed significant shape differences between all pairs of populations (*p* < 1e-04). The analysis of the angles between the divergence vectors suggests that both French populations diverged from their common ancestor in the same direction of the phenotypic space: these two vectors are more similar than two random vectors (α = 31.9°, *p* < 10^-4^, Fig. 2, Table S2). For the American populations, the angle between the two divergence vectors was greater than that of the French populations (α = 49.9°, *p* = 5 × 10^-4^, Fig 2, Table S2) and hint towards divergence along different directions. The average American and French direction of divergence from their common ancestor were strikingly different, and no more similar than random vectors (α = 71.9°, *p* = 0.11, Table S2). In terms of amount of divergence based on the Procrustes distance, the American populations appeared to have slightly less diverge than French populations, from their hypothetical ancestor (5.72 × 10^-3^, 7.46 × 10^-3^, respectively; Table S3) and as observed from the *bgPCA* (Fig. S2).

### The **G**-matrix for wing shape is mostly stable across populations

We found significant differences in the size of the **G** matrix among 3 pairs of populations, all involving the southern USA population US-WAT (Table 1). The **G** matrix of the US-WAT population had a significantly larger volume than those of US-DAY (northern USA) and of both French populations (FR-PAR and FR-MON). None of the other pairwise comparisons revealed any significant differences in the volumes of **G**. Regarding the shape of the **G** matrices, none of the **G** matrices exhibited pronounced eccentricity (i.e. no single PC dominated the others, inducing an elongated, “cigar-shaped” matrix). Visual inspection of the first two principal components indicated that all **G** matrices were approximately circular in two-dimensional space (with the exception of FR-MON), and would likely appear relatively spherical if represented as three-dimensional ellipsoids. (Fig. 3). Eccentricity values calculated as the ratio of the first to the second eigenvalues of **G** were indeed low and close to 1 for all matrices (Fig. S3). In other words, genetic variation in each population was relatively equally distributed among combinations of traits. We did not find any significant differences in the shape of the **G** matrices as eccentricity and the amount of variance explained by ***g****_max_* between any pairs of populations were not statistically different (Fig. S3). Similarly, no significant difference was found between the eigentensors of **S** calculated from the set of random and observed sets of **G** matrices based on the overlapping 95% HPD intervals (Fig. 4), suggesting a global stability of the **G** matrices across populations.

**Table 1.** Pairwise comparison of G matrices’ volumes. For each pair of **G** matrix the posterior mode of the ratio between volumes is shown along with the Highest Posterior Density (HPD) interval. Bold values correspond to significant differences between two matrices volumes whether the ratio is significantly > 1 (row matrix is larger) or < 1 (row matrix is smaller).

| Population | JP-TOK | JP-SAP | US-DAY | US-WAT | FR-PAR |
| --- | --- | --- | --- | --- | --- |
| JP-SAP | 1.03<br>[0.83:1.3] |  |  |  |  |
| US-DAY | 0.88<br>[0.69:1.08] | 0.87<br>[0.66:1.03] |  |  |  |
| US-WAT | 1.17<br>[0.95:1.45] | 1.14<br>[0.9:1.38] | **1.32**<br>**[1.09:1.65]** |  |  |
| FR-PAR | 0.93<br>[0.73:1.13] | 0.89<br>[0.7:1.07] | 1.05<br>[0.83:1.29] | **0.79**<br>**[0.63:0.94]** |  |
| FR-MON | 0.93<br>[0.73:1.11] | 0.88<br>[0.7:1.06] | 1.04<br>[0.83:1.26] | **0.76**<br>**[0.62:0.92]** | 0.95<br>[0.79:1.2] |

### Proportionality of **D** and ancestral **G**: a constrained response to selection?

Overall, the divergence matrix appeared fairly similar to the ancestral **G** matrix, its first two PCs and the ancestral **G** matrix respectively forming narrow angles of 13.6 and 15.8°, smaller than expected for random vectors (47.9°± 7.18). While the null hypothesis of proportionality between both matrices was not rejected (Bartlett-adjusted Χ² test; *p =* 1), suggesting a close match, the coefficient of proportionality (ρ = 1.063; 95% CI, 0.763 – 1.751) was in turn higher than the coefficient estimated for microsatellite markers (2F_ST_/(1-F_ST_) = 0.160; 95% CI, 0.128 – 0.198). Therefore, while **D** and **G** are proportional, their coefficient of proportionality deviates from the neutral expectation, suggesting a role for selection in the observed divergence.

### No reduction in genetic variance in the direction of divergence

We further projected each **G** matrix onto their respective direction of divergence from the hypothetical ancestor and estimated the corresponding percentage of variance (Fig. 5). Overall, we found that amounts of genetic variance in these directions were not statistically different among populations. Furthermore, we did not observe a lower amount of genetic variance between the projection of each population **G** matrix and the projection of the ancestral **G** matrix on their corresponding divergence vectors. This result goes against our expectation that selection would result in a depletion of genetic variance along each population’s divergence vector from the ancestor.

### The response to a theoretical selection does not match the observed divergences

To investigate the possible selective factors affecting wing shape evolution in *D. suzukii* invasion, we simulated the evolutionary responses to a hypothetical selection differential, the wing shape difference between fast and slow flying *D. suzukii* individuals, as reported in Fraimout *et al*. (2018; Fig. 6). The evolutionary response is shown on Fig. 6, as the average response derived from the full posterior of MCMC samples. The shape divergence of French and American populations from their common hypothetical ancestor are shown on Figure 6c and d (the two Japanese populations used to compute the ancestral **G** matrix were considered too close from this ancestral state and were thus not considered). The pattern of shape change associated with ***S_velocity_*** and the average response *Δ**z*** were very similar (mean α = 22.74°, SD 6.03) confirming that **G** does not constrain the response to selection. This result is in line with the relatively spherical shape of the ancestral **G** matrix, which should not constrain the response to selection. While a similarity in the shift of the basal landmarks can be found between the divergence of French populations and the theoretical response to selection, no obvious similarity with American populations divergences is detected (Fig 6, Fig. S4). However, this congruence was not reflected in the correlation between vectors when accounting for the full landmark configurations as the evolution of the distal part of the wing did not match the expected response to selection (Fig. 6, Fig. S4). Overall, the similarity between the divergence vectors from each population and the response to selection *Δ**z*** was low (Fig. S4) suggesting that the observed divergence in wing shape does not result from selection on faster flight capacity.

**Figure 6.**
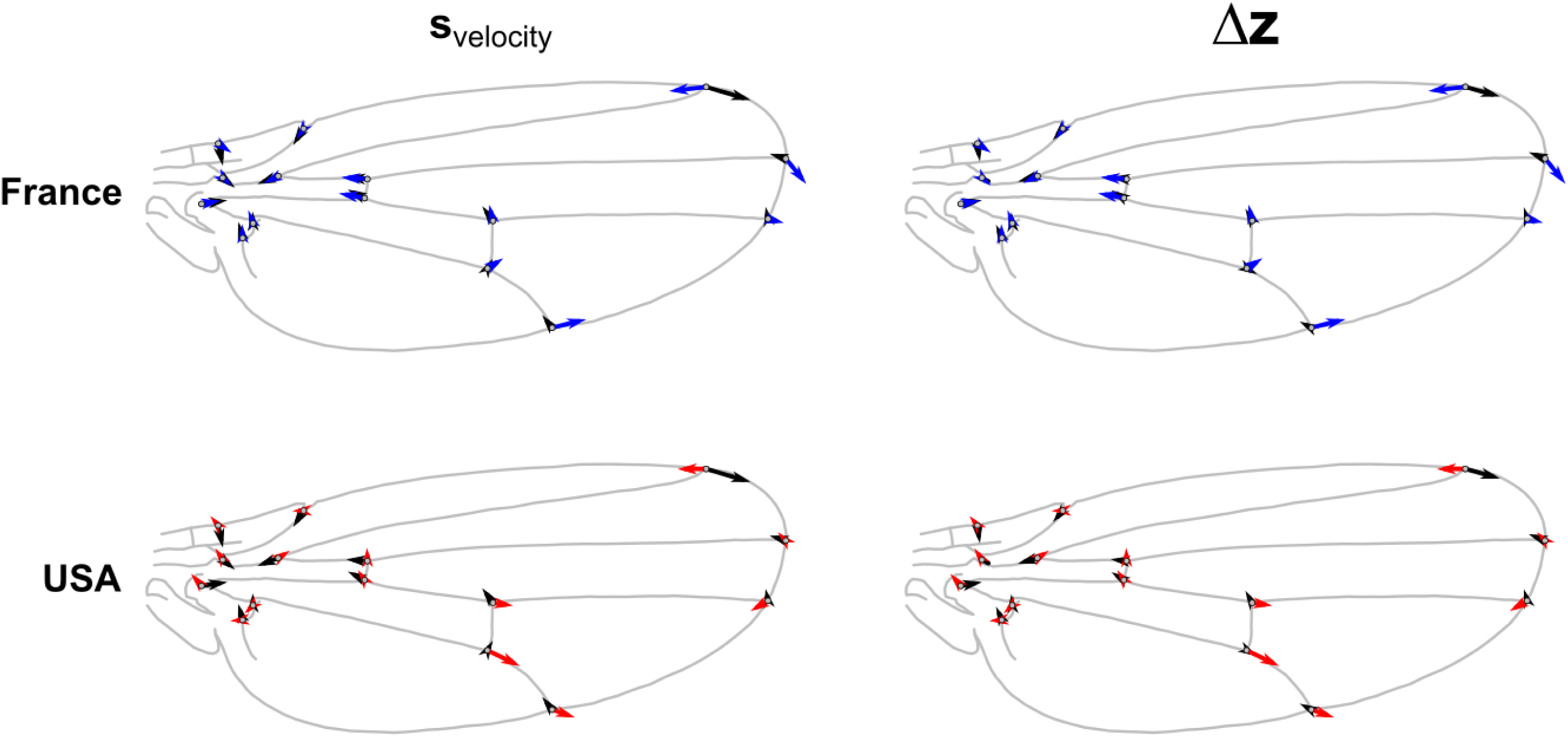
Response to a hypothetical selection compared to observed shape changes. The observed wing shape change of the average French (blue arrows) and USA (red arrows) population are shown, representing the directions of change for each landmark, from the ancestral wing shape. The shape change associated with the selection differential defined by the shape difference between slow and fast fliers from Fraimout et al (2018) (**s**_velocity_) is shown with black arrows on the left panels. The shape change corresponding to the response to selection (Δz) based on the MCMC estimates of the ancestral **G** and **P** matrices is shown with black arrows on right panels. The results show similarity in the shift of the basal landmarks between the divergence of French populations and the theoretical response to selection (alignment between blue and black arrows of the leftmost landmarks; upper right panel) but no congruence in any other comparison, suggesting limited evidence for selection on flight performance in these populations.

## Discussion

### Genetic divergence of wing shape across populations of Drosophila suzukii

Geographical variation of wing morphology has been described in several *Drosophila* species (*e.g.* Azevedo *et al*., 1998; Haas & Tolley, 1998; Hoffmann & Shirriffs, 2002), including introduced ones (Huey *et al*., 2000). In particular, geographical clines of wing size and shape have been detected along latitudinal and altitudinal gradients (Huey *et al*. 2000; Gilchrist *et al*., 2001; Gilchrist *et al*., 2004; Pitchers *et al*., 2013; Adrion *et al*., 2015). These clines have typically been interpreted as adaptations to flight conditions, specifically temperature and air pressure (Dillon & Frazier 2006; Pitchers *et al*. 2013), although direct evidence on the nature of selection is lacking. In *D. suzukii*, clines in wing morphology have been reported along an altitudinal gradient in Hawaii – the first and oldest invaded territory (Asplen *et al*., 2015) – with flies from high altitude (ca. 2000m) displaying significantly larger wings than flies sampled at sea level (Koch *et al*., 2020). Here, we detected geographic variation of wing shape across all populations, but the differences among north and south populations were different among continents. Gilchrist *et al*. (2000) reported differences in latitudinal wing shape changes in *D. melanogaster* from three continents, and suggested the occurrence of specific local selective pressures on wing shape. However, they also detected parallel clines in wing size across continents, suggesting common adaptive effects. In our study, wing size was more stable: the only difference between northern and southern populations was found in the USA. As for wing shape, although we found that populations from the same continent have evolved in a relatively similar direction of the phenotypic space, we did not find any evidence for parallel latitudinal clines. At the time of sampling, the invasion of *D. suzukii* was initiated less than a decade ago in Europe and the USA and it is possible that adaptive geographical clines did not have enough time to evolve in these regions, contrary to the first invaded range in Hawaii (Koch *et al*. 2020). As a comparison, a latitudinal cline in *D. subobscura* following colonization of the New World, was reported after two decades, while no difference had been detected ten years after the start of the invasion (Huey *et al*., 2000; Gilchrist *et al*., 2004). At this point, it is unclear whether *D. suzukii* wing phenotypes are locally adapted, and additional work is warranted to decipher the interplay between wing plasticity, genetics and their association with fitness in natural *D. suzukii* populations. Nevertheless, by rearing flies in controlled laboratory conditions for a few generations, we showed that the phenotypic divergence observed among populations is genetic rather than plastic (see also Fraimout *et al* 2018) and that some evolutionary process independent of phenotypic plasticity have shaped the wings of *D. suzukii* in the wild.

### **G** matrix of wing shape is stable across populations

Despite the wealth of theoretical and empirical research (e.g. Roff, 2000; Steppan *et al*., 2002; Bégin & Roff, 2003; Jones *et al*., 2003; Mezey & Houle, 2003; Björklund, 2004; Jones *et al*., 2004; Blows & Hoffmann, 2005; Arnold *et al*., 2008; Walsh & Blows, 2009; Jones *et al*. 2014; Wood & Brodie, 2015; Puentes *et al*., 2016; Svensson *et al*., 2021), predicting how natural selection or drift will affect the structure of **G** and at which pace **G** can evolve in the wild remains a difficult task. As stated by Jones *et al*. (2003): “[…] *the current state of analytical theory on **G**-matrix evolution is that existing theory cannot guarantee stability of the **G**-matrix (Turelli 1988), but also does not guarantee instability*.” Following this assertion, simulation studies (Jones *et al*., 2003; Jones *et al*., 2004; Arnold *et al*., 2008; Jones *et al*., 2014; Björklund & Gustafsson, 2015; Chantepie & Chevin 2020) have identified how correlative selection, pleiotropic mutations, population size and other evolutionary forces can affect the stability of **G**’s structure and orientation. However, empirically and 20 years later, evidence for both stability (Delahaie *et al*., 2017; McGlothlin *et al*., 2018, Henry & Stinchcombe 2023) and lability (*e.g*., Uesugi *et al*., 2017, Chantepie *et al*. 2024) have been reported in varying contexts.

Here we found that the **G** matrices were globally stable across populations, the differences between populations mainly pertaining to the volume of **G** (i.e. the global amount of genetic variance). Specifically, French populations seemed to display less genetic variance as indicated by their smaller volumes, while the US-WAT population seemed more genetically variable than the rest of the populations. Reduction of genetic variance and, therefore, of the volume of **G**, is expected under certain conditions such as increased genetic drift due to reduced effective population size (Shaw *et al*., 1995; Roff, 2000; Jones *et al*., 2004) or increased strength of stabilizing selection (Jones *et al*. 2003). According to the results of Fraimout *et al*. (2017) obtained from neutral molecular markers and Olazcuaga *et al*. (2020) using whole-genome data, all invasive populations of *D. suzukii* experienced significant loss of genetic diversity due to various intensities of bottlenecks upon introduction. Lesser levels of genetic variance and lower volumes in the French **G** matrices are thus consistent with the effect of drift. Similarly, the higher volume of the US-WAT population could be explained by the genetic admixture of two highly differentiated lineages from Asia and Hawaii (Fraimout *et al*. 2017). However, populations’ invasion history was not reflected in all **G** matrices, as other genetically admixed populations (FR-PAR, US-DAY; Fraimout *et al*. 2017) did not show higher volumes and neither did ancestral populations where genetic bottlenecks most likely did not occur. Contrasting with the random evolution of **G** in all directions of the multivariate space expected under genetic drift (Arnold *et al*., 2001; Phillips *et al*., 2001), our results show that most aspects of the eigenstructure of **G** (size and shape) have remained stable throughout the invasion process.

Our analyses may lack statistical power to detect more subtle differences between **G** matrices. Indeed, while our crossing design consists of an acceptable number of genetic lines per population, the multidimensional nature of the *Drosophila* wing implies a large number of parameters to be estimated (n x (n-1) /2 + n; 351 in our case), larger than the number of replicates (*i.e.* isofemale lines). Nonetheless, replicating our analyses using fewer numbers of PCs - and thus less parameters to be estimated - led to the same conclusions (see *Supplementary material*). Finally, our design did not allow us to partition additive from non-additive genetic components of wing shape variation. Dominance genetic variance has been shown to explain up to 35% of phenotypic variation for wing traits in *D. serrata* (Sztepanacz & Blows 2015) and it is therefore possible that some of our estimates of genetic variance could be inflated by such non-additive components. Furthermore, the use of a limited number of isofemale lines to capture the possibly large standing genetic variation found in *D. suzukii*’s wild populations could further exacerbate the effects of non-additivity on our estimates of the **G** matrix. Specifically, establishing isofemale lines could result in inbreeding depression, where mildly deleterious mutations otherwise sheltered in heterozygotes are expressed in the laboratory under sib-mating (Agrawal & Whitlock 2012). This in turns could inflate the contribution of dominance genetic variance (Charlesworth & Willis 2009). While we did not estimate the contribution of dominance variance, the use of gravid females mated in the wild may have reduced the effect of inbreeding depression upon establishing isofemale lines. Whether between-population differences exist in the importance of dominance variance for wing shape is unknown, and future work including appropriate breeding designs (*e.g.,* double-cousins) should shed light on the importance of non-additive genetic components in wing shape variation of *D. suzukii*.

### Phenotypic divergence is not constrained in D. suzukii’s invasive populations

We tested the hypotheses of ancestral constraints on wing shape evolution by estimating the similarity between the ancestral **G** matrix and the divergence matrix **D**, and found that the leading vectors of both matrices were tightly associated. Such similarity can be viewed as a mark of genetic constraints affecting divergence, compatible with both neutral and adaptive scenarios. Genetic drift is expected to induce divergence along the axes with the most genetic variation, and proportionally to the amount of variation in the ancestral population (Lande 1979; Lofsvold 1998; Marroig & Cheverud, 2004). But this pattern of concordance between **G** and **D** is also expected under selection, the main directions of genetic variation biasing the response to selection (i.e. the lines of genetic least resistance; Schluter, 1996). However, the estimated ancestral **G** matrix for *D. suzukii* wing shape had a relatively spherical shape, suggesting the existence of genetic variation in all directions of the phenotypic space, and therefore a lack of genetic constraint. This is evident from the results of our phenotypic divergence vector analysis showing that, despite the relatively small differentiation among populations, the evolution of wing shape in France and USA has followed very different directions (Fig. 2), suggesting no constraint. Finally, the response to selection predicted from our theoretical selection gradient applied to the estimated ancestral **G** produced a wing shape different from that observed in any of the extant invasive populations, further supporting an absence of genetic constraint on wing shape evolution in this species. These results are interesting in the light of recent research investigating the link between trait evolvability and divergence (Tsuboi *et al*. 2024, Pélabon *et al*. 2025). Recent studies have shown that population divergence across species and traits tends to scale positively with standing additive genetic variance (*i.e.,* evolvability; Opedal *et al*. 2023, Holstad *et al*. 2024), suggesting that phenotypic evolution can be channeled by ancestral genetic constraints through lines of least resistance. Here, we did not find evidence for genetic lines of least resistance, but rather, unconstrained phenotypic divergence from ancestral standing genetic variation.

### Adaptive or neutral divergence?

The results from our multivariate Q_ST_-F_ST_ support an adaptive scenario for wing shape evolution, as divergence was found to be significantly greater than expected under neutral evolution. While failure to account for demographic history and population structure can bias the outcome of Q_ST_-F_ST_ comparisons (de Villemereuil *et al*. 2022, do O *et al*. 2025), it does not seem to be the case here. The demographic history of *D. suzukii* is known, and there is no evidence for gene flow or migration among populations (even within-country; Fraimout *et al*. 2017) that could reduce values of F_ST_, biasing upward the result for our test of selection (ρst > F_ST_). As for a possible inflation of ρst, this could happen if all populations had undergone drastic bottlenecks and reductions in **G** while **D** remained large. We do not have evidence that this happened in our study populations, as the volume of most invasive populations’ **G** is not significantly smaller than that of the native population. The divergence in wing shape among invasive populations of *D. suzukii* might thus be adaptive. The statistical power of this test is conditional on the sampling design and particularly the number of populations sampled (Martin *et al*. 2008). While the test for departure of neutrality should be robust to a small number of populations (*i.e.,* < 10 as in the current study) results of the proportionality test should be interpreted with caution. Similar analyses using long-diverged groups (*i.e.,* Teosinte and Maize), report values of ρ_ST_ several order of magnitude above the neutral expectation (Yang *et al*. 2019), suggesting that the signal of selection we observe in *D. suzukii* is rather low. The adaptive value of wing shape variation in *Drosophila* is a long-standing conundrum, and in the absence of fitness data, remains speculative (Gilchrist *et al*. 2000). Given the role of wing in flight, we leveraged available flight data (Fraimout *et al* 2018) to test whether selection on flight speed could be involved in wing shape divergence among our populations. Despite partial correspondence, the response to this selection applied to the ancestral **G** matrix, was very different from the observed evolutionary divergence, providing no support for selection on flight speed as a driver of this divergence.

Finally, it is worth stressing that we focused our analyses on the study of male wings only. Wing shape is under strong sexual selection in Drosophila, as males produce a variety of visual and acoustic signalling using their wings during courtship (Debelle *et al*. 2014). By focusing on male wings only, it is possible that we are missing an important component of evolutionary constraint, as previous work has shown that genetic covariance across sexes can bias the evolutionary response of wing shape to selection (McGuigan & Blows 2007, Sztepanacz & Houle 2019). Nonetheless, recent work on *D. buzzati* (Iglesias *et al*. 2023) suggests that adaptive divergence in wing shape can be attributable to selection on courtship traits in males among populations. Additionally, Branden & De Lisle (2026) have shown that the effect of stabilizing selection is particularly pronounced in male Drosophila, where departure from the average wing shape is associated with lower fitness in males. Our results could be compatible with those studies, reflecting a combination of stabilizing selection on *D. suzukii* males’ wing shape and divergent selection among populations. Without access to fitness assays and its relationship to wing shape variance in our study, this hypothesis cannot however be formally tested with the data at hand. The nature of the selective factors affecting wing shape evolution thus remains to be identified in *D. suzukii*, and further work would be valuable to shed light on the relationship between wing shape and fitness in males, and on the effect of stabilizing selection on the wing shape **G** matrix.

### Conclusion

Consensual predictions on the effect of selection and drift on the **G** matrix remain scarce in the literature and most empirical evidence are based on experimental evolution studies (*e.g.* Doroszuk *et al*., 2008; Careau *et al*., 2015) or comparisons among long-diverged species (e.g. McGlothlin *et al*., 2018). In this context our study provides new insights on the evolution (or lack thereof) of **G** over a relatively short evolutionary time scale in natural conditions. We showed that native and invasive populations of *D. suzukii* display quantitative genetic differentiation in wing shape across continents and at an early stage of their worldwide invasion. Our results suggest an effect of selection acting on a poorly constraining ancestral G matrix. They provide a reference point for future studies quantifying phenotypic evolution in this species.

## Acknowledgments

AF was supported by the Agence Nationale de la Recherche, through the LabEx ANR-10-LABX-0003-BCDiv, of the program “Investissements d’avenir” (ANR-11-IDEX-0004-02). VD was funded by ANR SWING (ANR-16-CE02-0015). The authors are grateful to S. Fellous, A. Xuéreb, K. Tamura, M. Toda, P. Shearer, T. Schlenke and A. Kopp for help with flies sampling. We thank M. Guillaume for helping with the stock flies maintenance and F. Peronnet for providing flies rearing medium.

## Author contribution

AF, VD and CT designed the study; AF, SC and NN analyzed the data; AF and VD obtained the samples; AF led the writing of the first draft and all authors contributed critically to the manuscript.

## Data availability

Data and code necessary to replicate the analyses presented in the manuscript have been deposited on the Zenodo platform (zenodo.org).

