## Supplementary material for "Wing shape evolution is not constrained by ancestral genetic covariances in the invasive *Drosophila suzukii*"

**Short title:** Quantitative genetics of *Drosophila suzukii* wing shape

[1]Institut de Systématique, Evolution, Biodiversité, ISYEB – CNRS, MNHN, UPMC, EPHE, Université des Antilles, Muséum National d'Histoire Naturelle, Sorbonne Universités, 45 rue Buffon, 75005 Paris, France
[3]UMR CNRS 6282 Biogéosciences, Université de Bourgogne-Franche-Comté, 21000 Dijon, France
[4]Centre d'Ecologie Fonctionnelle et Evolutive, CEFE – UMR 5175 – CNRS, IRD, EPHE, Université de Montpellier, Université Paul Valéry, 34293 Montpellier, France

*These authors contributed equally to this work

### TABLE OF CONTENTS


| | |
|---|---|
| **Figure S1. Population differences in wing centroid size** | Page 2 |
| **Figure S2. Results of the Between-group Principal Component Analysis on wing shape** | Page 3 |
| **Figure S3. Between population comparison of gmax and eccentricity.** | Page 4 |
| **Fig. S4. Angles between divergence vectors and the response to selection.** | Page 5 |
| **Table S1. Summary of the population samples.** | Page 6 |
| **Table S2. Angles between the divergence vectors from the ancestral G matrix** | Page 7 |
| **Table S3. Distance from the hypothetical ancestor** | Page 8 |

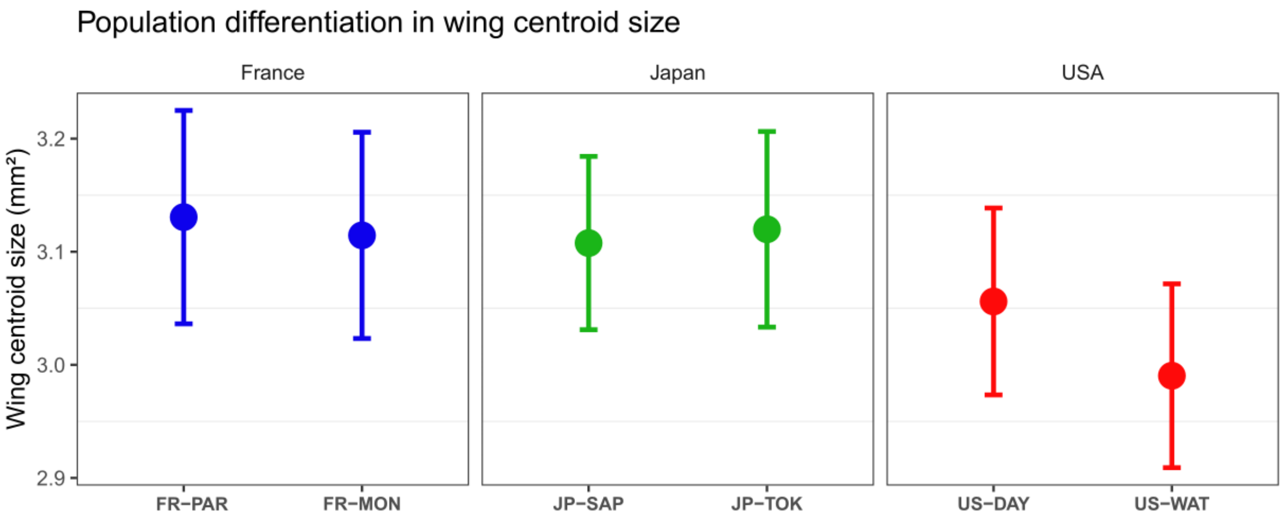


**Figure S1. Population differences in wing centroid size.** Mean values of wing centroid size are shown for each population and colored according to the country of origin as in Fig. 1 (green = Japan, blue = France, red = USA). Vertical error bars represent standard deviation.

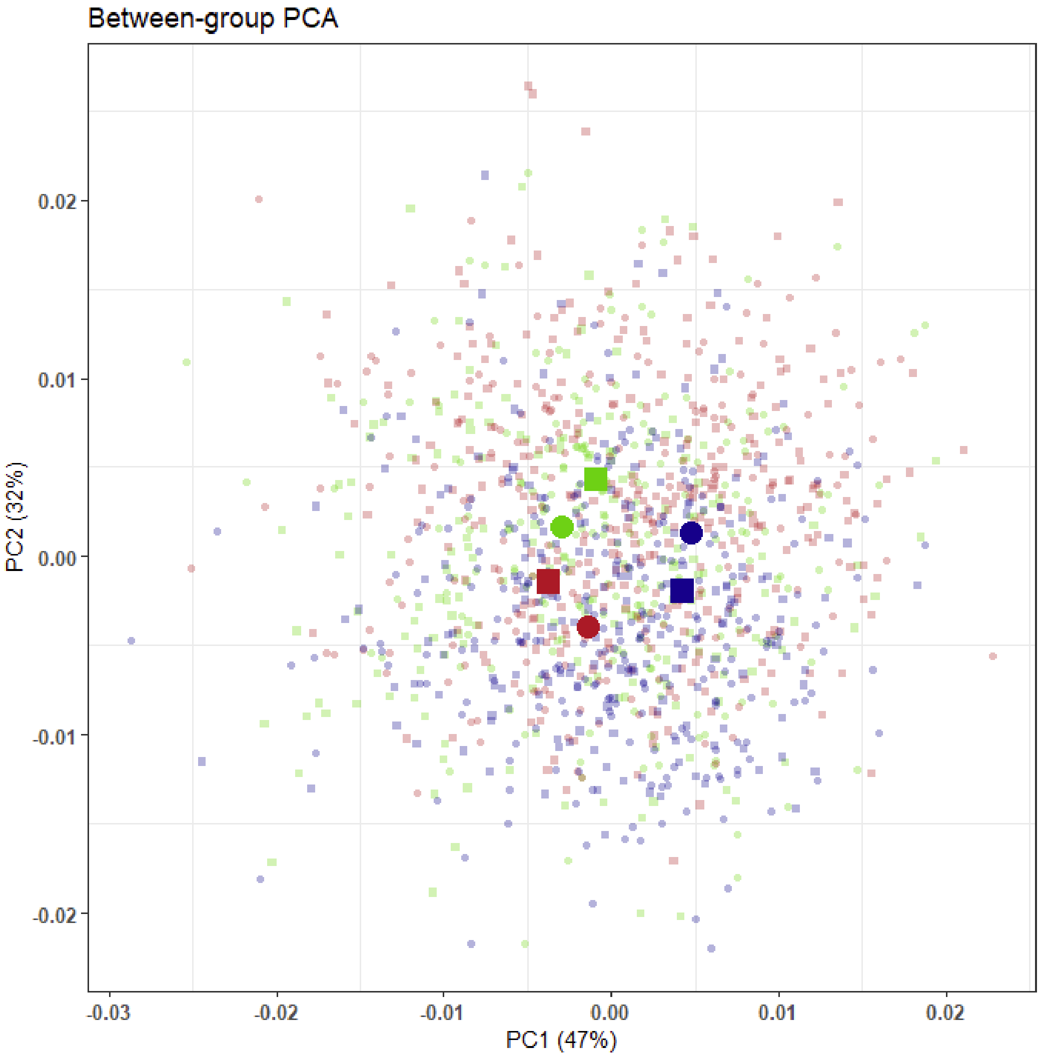


**Figure S2. Results of the Between-group Principal Component Analysis on wing shape.** Individuals are colored according to their country of origin as in Fig. 1 (green = Japan, blue = France, red = USA) and shape represent the latitude of the population of origin (square = North, circle = South).

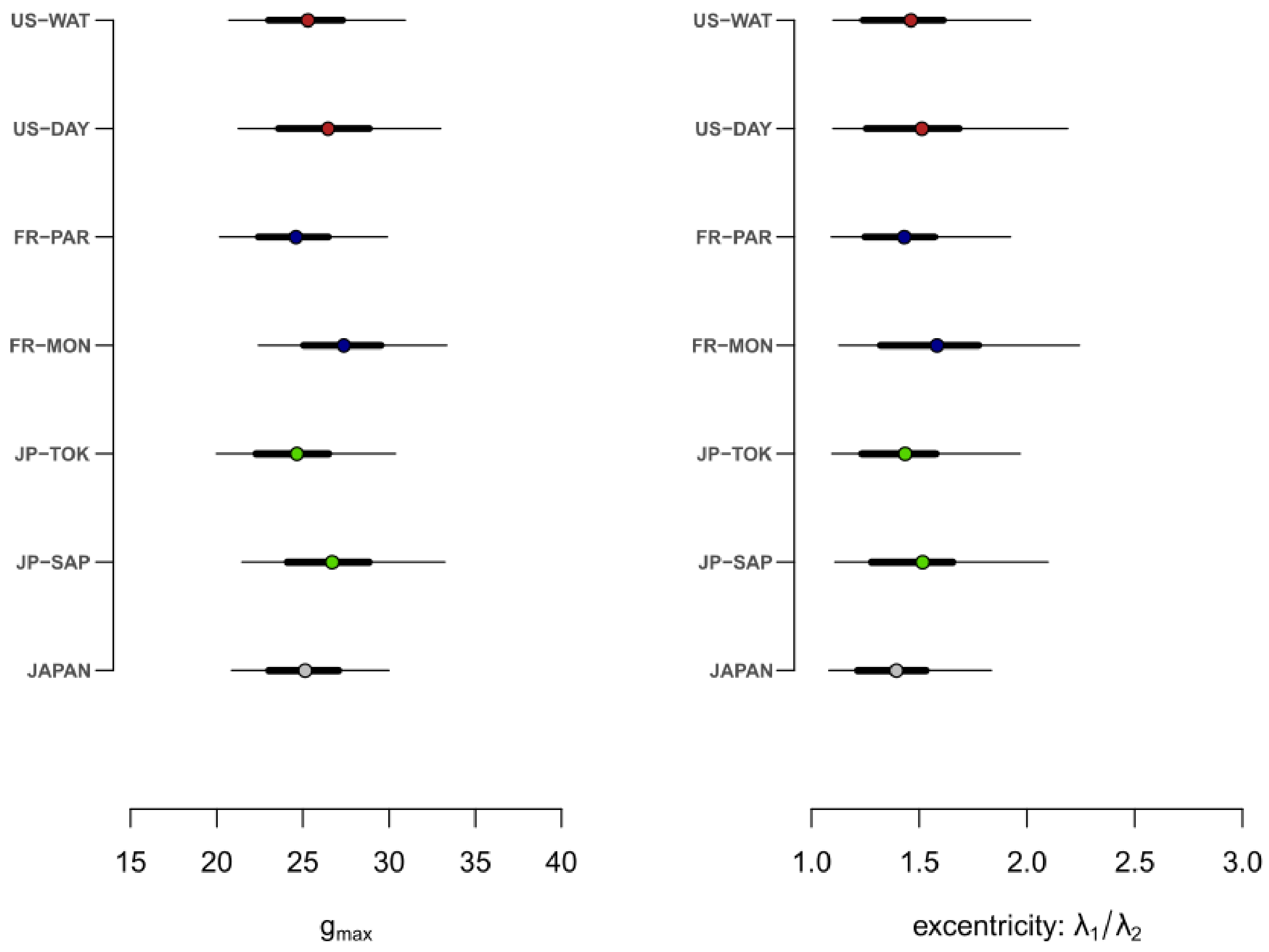


**Figure S3. Between population comparison of gmax and eccentricity.** The amount of genetic variance along the leading eigenvector (gmax, left panel) of each population **G**-matrix (Population code) and for the hypothetical ancestor (JAPAN) is shown (filled circles) with the 95% Highest Posterior Density interval (black horizontal lines). Eccentricity (shape of the matrix) of each population **G**-matrix and of the ancestral **G** is shown (filled circles) with the 95% Highest Posterior Density interval (black horizontal lines). In both panels, overlapping of confidence intervals indicates a lack of statistically significant difference. Colors correspond to the sampled populations as in Fig. 1.

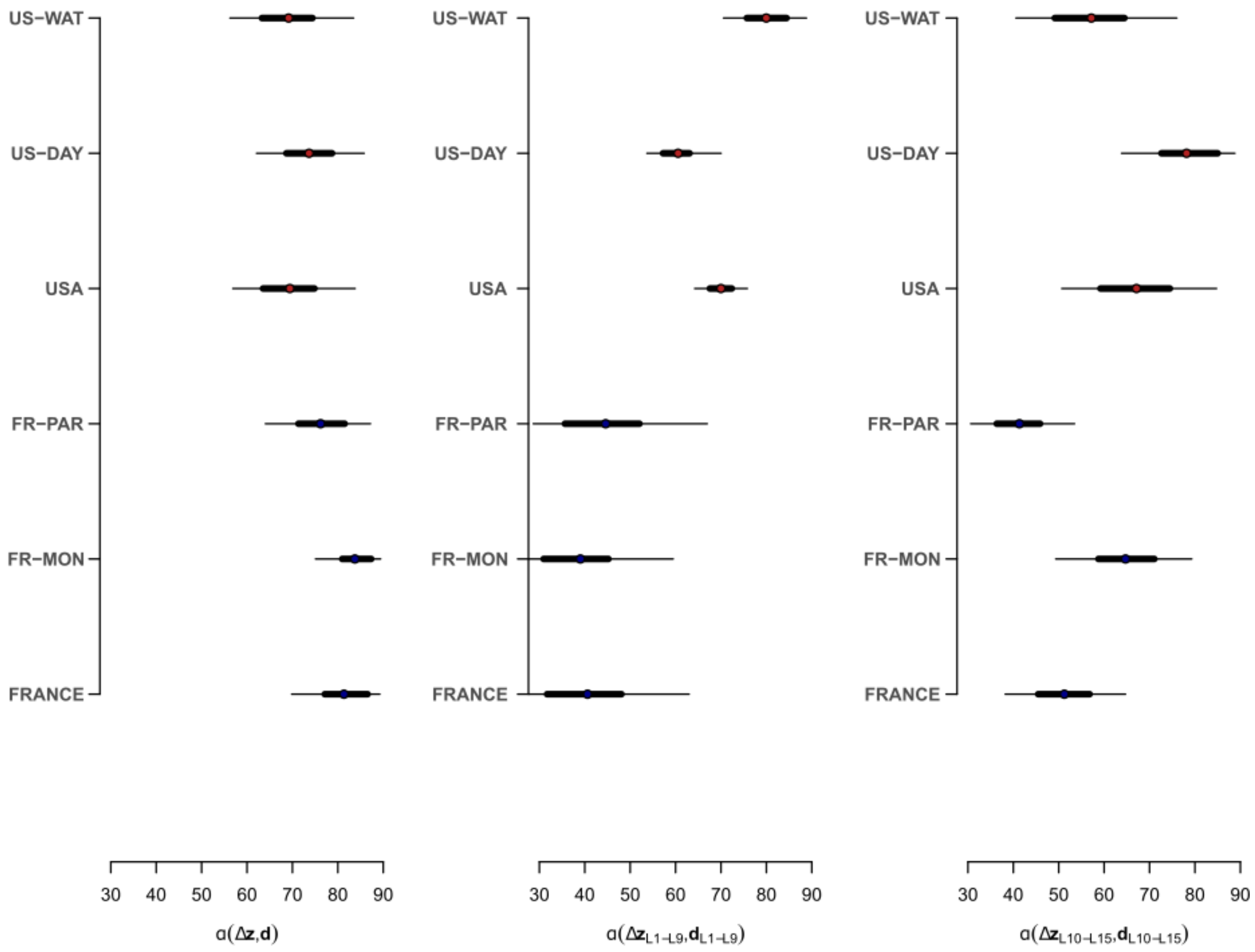


**Fig. S4. Angles between divergence vectors of the invasive populations and the response to selection.** The angle (α) between each invasive populations or the average invasive population's divergence vector from the ancestor (d) and the response to selection (Δz) is shown (filled circles) with the 95% Highest Posterior Density interval (black horizontal lines) based on the whole landmark configuration (left panel); for the basal landmarks only (middle panel) and for the distal landmarks (right panel). Colors correspond to the country of origin as in Fig. 1.

**Table S1. Summary of the population samples.** For each population the table indicates the sampling location (city) within the corresponding country and the associated population code used throughout the manuscript. The column "Status" indicates whether the population was sampled in the native or the invasive range. Sample sizes (N) are shown for the number of individuals and the number of isofemale lines (in brackets) per population

| Sampling location | Country | Status | Population code | N |
|---|---|---|---|---|
| Tokyo | Japan | Native | JP-TOK | 192 (31) |
| Sapporo | | | JP-SAP | 192 (24) |
| Dayton | USA | Invasive | US-DAY | 313 (38) |
| Watsonville | | | US-WAT | 191 (26) |
| Paris | France | | FR-PAR | 209 (25) |
| Montpellier | | | FR-MON | 226 (33) |

**Table S2. Angles between the divergence vectors from the ancestral mean wing shape**. Above diagonal elements show the angles between each pairwise combination of divergence vectors from the hypothetical ancestral Japanese population mean. Below diagonal show the *p*-value for the significance of each pairwise comparison compared with the differences with 1000 random vectors (see *Methods*) with bold values indicating statistical significance ($p < 0.05$).

| | FR-MON | FR-PAR | US-DAY | US-WAT | USA | FRA |
|---|---|---|---|---|---|---|
| FR-MON | | 31.929 | 85.758 | 75.513 | 79.699 | 16.783 |
| FR-PAR | **< 0.001** | | 80.570 | 55.120 | 66.055 | 15.146 |
| US-DAY | *0.711* | *0.418* | | 49.947 | 24.982 | 82.762 |
| US-WAT | *0.212* | **0.002** | **0.001** | | 24.964 | 64.164 |
| USA | *0.368* | **0.035** | **< 0.001** | **< 0.001** | | 71.945 |
| FRA | **< 0.001** | **< 0.001** | *0.546* | **0.024** | *0.119* | |

**Table S3. Distance from the hypothetical ancestor.** The Procrustes distance from the hypothetical Japanese ancestor (see *Methods*) is shown for each population separately and for the combined French (France) and North American (USA) populations.

| Population | Distance |
|---|---|
| JP-TOK | 2.754 |
| JP-SAP | 2.754 |
| US-DAY | 6.311 |
| US-WAT | 6.315 |
| FR-PAR | 8.151 |
| FR-MON | 7.376 |
| France | 7.465 |
| USA | 5.723 |
